# Drug-Target Binding Affinity Prediction Using Transformers

**DOI:** 10.1101/2021.09.30.462610

**Authors:** Mahsa Saadat, Armin Behjati, Fatemeh Zare-Mirakabad, Sajjad Gharaghani

## Abstract

Drug discovery is generally difficult, expensive, and low success rate. One of the essential steps in the early stages of drug discovery and drug repurposing is identifying drug-target interactions. Binding affinity indicates the strength of drug-target pair interactions. In this regard, several computational methods have been developed to predict the drug-target binding affinity, and the input representation of these models has been shown to be very effective in improving accuracy. Although the recent models predict binding affinity more accurate than the first ones, they need the structure of target proteins. Despite the strong interest in protein structure, there is a massive gap between known sequences and experimentally determined structures. Therefore, finding an appropriate presentation for drug and protein sequences is vital for drug-target binding affinity prediction. In this paper, our primary goal is to assess the drug and protein sequence representation for improving drug-target binding affinity prediction.

## I. Introduction

Drugs are often developed to target proteins involved in many cellular processes. Thus, identifying drug-target interactions (DTIs) has become an essential step in the early stages of drug discovery. Most DTI prediction computational methods focus on a binary classification to find if the interaction between a drug and its target exists[1]. However, there are more challenges in predicting the binding strength (affinity) between a drug and its target[2].

Binding affinity indicates the strength of drug-target pair interactions. Through binding, drugs can positively or negatively affect the functions performed by proteins which eventually leads to the treatment of the disease. Since the high binding affinity between a small molecule and a target protein is one of the criteria for selecting a candidate compound in drug discovery[3], the drug-target binding affinity (DTBA) discovery has received much attention in recent years. The time and financial expenses of directly measuring the binding affinity through experimental methods are incredibly high. Therefore, there is a need to develop computational models for binding affinity prediction. So far, several computational models have been developed to predict the binding affinity of drug-target pairs[4]. DTBA prediction models often take drug and protein data as inputs to compute binding affinity as a regression problem.

These models can be divided into two categories based on types of feature extraction from the input (Fig. 1): manual feature extraction and automatic feature extraction. The first two machine learning models introduced for this problem are called KronRLS[2] and SimBoost[5], which follow the “guilt by association” rule. This rule is based on the assumption that similar drugs tend to interact with similar targets, and similar targets are interacted by similar drugs[4]. These two models manually extract the features of target and drug based on similarities. Manually appropriate feature extraction requires a great deal of biological pre-knowledge[4]. In addition, some of these features may be highly correlated and lead to data redundancy. In this regard, automatic feature extraction methods such as auto-encoder, transformer are developed using deep learning to overcome these challenges[6].

**Fig. 1.**
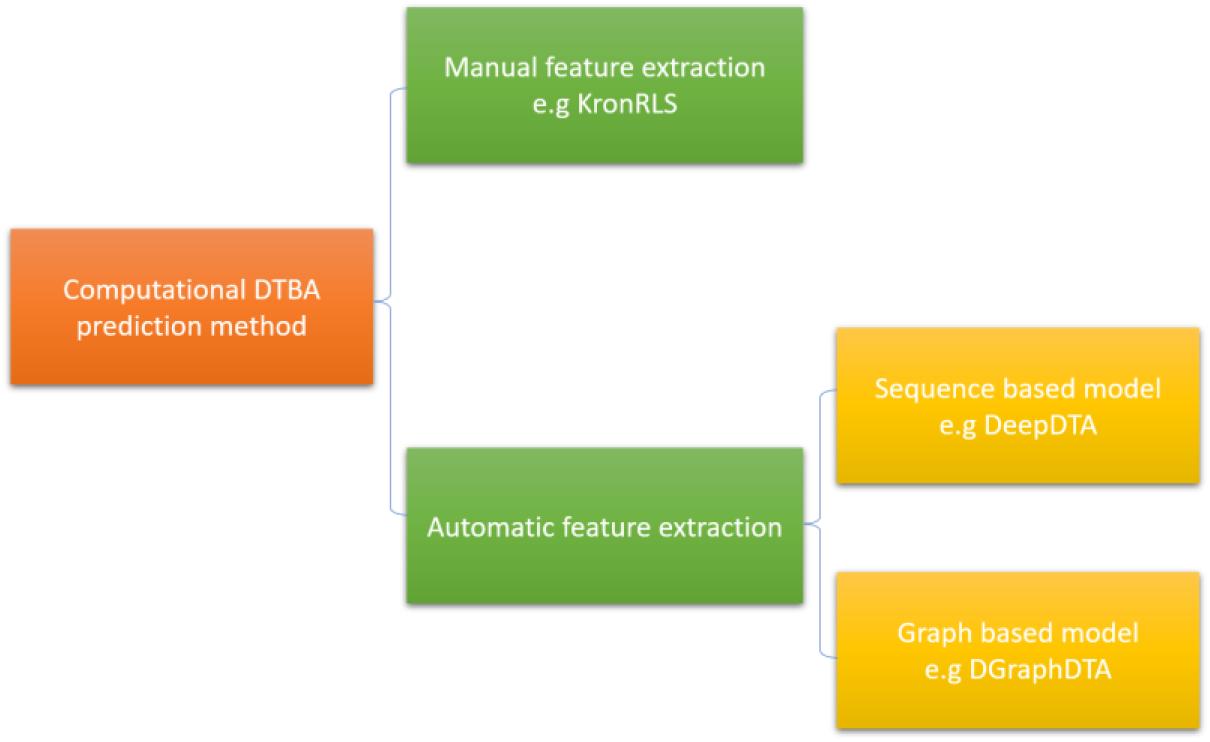
Computational DTBA prediction methods.

Automatic feature extraction has been categorized into two classes, sequence-based and graph-based models. In 2019, DeepDTA[7] was introduced as the first sequence-based deep learning model for predicting DTBA. Drug simplified molecular-input-line-entry system (SMILES) and protein sequence are used as inputs in this model. Each character of the sequence and SMILES is encoded by an integer number [7]. So, there are two integer vectors for protein sequence and drug SMILES, respectively. For feature extraction, these vectors are given to two convolutional neural networks (CNN) blocks. Then protein and drug features are concatenated to feed fully connected neural layers for binding affinity prediction[7]. Although DeepDTA[7] performs better than KronRLS[2] and SimBoost[5], this model use sequences as inputs while the useful biological information for DTBA lies in drug and target structures. It seems from the fact that the structure of the molecule is more applicable in the binding affinity concept[8]. In this regard, some graph-based models such as GraphDTA[9] and DGrapgDTA[10] are introduced to use the drug and protein structures as inputs.

Although deep learning models show excellent performance improvement in DTBA prediction, two main challenges remain to study. First, using structural information in these models increases the cost of time and resources. Second, the 3D structure is not available in most cases, so the structure needs to be predicted. The use of predicted information backpropagates the error rate on the main issue.

Therefore, finding an appropriate presentation for protein and drugs to extract more information from the sequences for drug-target binding affinity prediction can be an important step in scientists’ research. Recently, transformers have been known as models to extract structural information from drug and protein sequences[11], [12]. Also, recurrent neural network (RNN) based models such as UniRep[13] are used for protein sequence feature extraction.

In this paper, our primary goal is to assess protein sequence representation with ProtAlbert[12], ProtBert[12] and UniRep[13] models, and also drug representation with RoBERTA[14] and PubChem[15] fingerprint. We concatenate two extracted vectors from the model representations of the protein and drug s to feed a multi-layer fully connected neural network to predict DTBA. We perform the model on KIBA[16], a benchmark dataset for binding affinity prediction evaluation[2].

The best result is obtained by ProtAlbert[12] for protein representation and molecular fingerprint representation for the drug. The architecture of ProtAlbert[12] is based on the ALBERT transformer[17]. It is known as one of the best pre-training transformers on protein sequences, which their layers and heads are protein structural representative[18]. The results show that our proposed method, TranDTA, achieves a competitive performance in the DTBA prediction without relying on structure information.

## II. Materials and Methods

We formulate the DTBA prediction as a regression task to determine how strong a pair of drug and target protein bind. In DTBA, protein *t* and drug *d* are given as inputs. A real value is generated as the output to show the binding affinity between target *t* and drug *d*. In this section, we introduce input representations and our proposed model.

### A. Input representation

In this section, we introduce different representations of the inputs.

#### 1) Drug representation

In the following, two representations are explained for each drug.

- **Molecular Fingerprint:** We use PubChem molecular fingerprint[19] representations to represent drug inputs. In cheminformatics, molecular fingerprints are one of the most common representations of chemical structures. This representation increases the calculation speed and reduces the storage space[20]. The fingerprint representation of each drug *d* is shown by an 881-length vector, as follows:

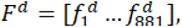

where 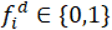 indicates the absence (0) or presence (1) of a substructure descriptor associated with a specific molecular feature predetermined in the design of the fingerprint.
- **SMILES representation by RoBERTa:** Using the SMILES sequence of each drug can be found in PubChem[15]. We convert the SMILES sequence to a numerical vector by RoBERTa. The RoBERTa[14] model is an extended model of BERT[21] with a change in its pre-training process. It is a transformer-based language model with 12 layers and 12 attention heads. This study uses pre-trained RoBERTa with 541,555 SMILES sequences[22]. In RoBERTa representation, each drug such as *d* is shown by a 768-length vector, as follows:

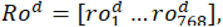

where 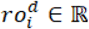 and 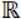 is the real number set.

#### 2) Protein representation

In the following, we introduce the representations of the target protein. Each sequence of target protein *t* is defined as follows:

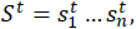

where 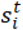 shows one out of 20 amino acid types, and *n* is the length of the protein sequence. Here, we encode protein sequence as a vector with different models:

- **UniRep:** UniRep[13] is a pre-trained model that uses a multiplicative long short term memory (mLSTM) to learn protein representation. Without using structural data, UniRep approximates the protein sequence to a constant length vector containing the protein’s fundamental (structural and amino acid) properties. In UniRep, each target protein *t* is represented as a 1900-length vector, as follows:

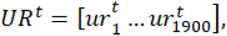

where 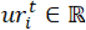.

It has recently been shown that encoded protein sequence representations from transformers-based models can capture the biophysical and structural features of the original sequence[11]. We use ProtBert and ProtAlbert as protein sequence encoders[12]. They were trained on the Uniref100 dataset[23] that includes 216 million protein sequences.

- **ProtBert:** ProtBert[12] is a transformer-based language model with 30 layers and 16 attention heads. It generates a 1024-length vector for each protein sequence as follows:

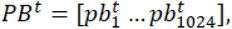

where 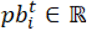.
- **ProtAlbert:** ProtAlbert[12] is a transformer-based language model with 12 layers and 64 attention heads. It generates a 4096-length vector for each protein sequence as follows:

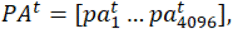

where 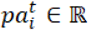.

### B. Model

The DTBA problem is considered a regression model. We propose a framework that feeds the concatenated protein and drug representations to a multi-layer fully connected neural network. This framework leads us to find the best representation for drugs and proteins based on sequences. Fig. 2 shows the step-by-step of our framework.

**Fig. 2.**
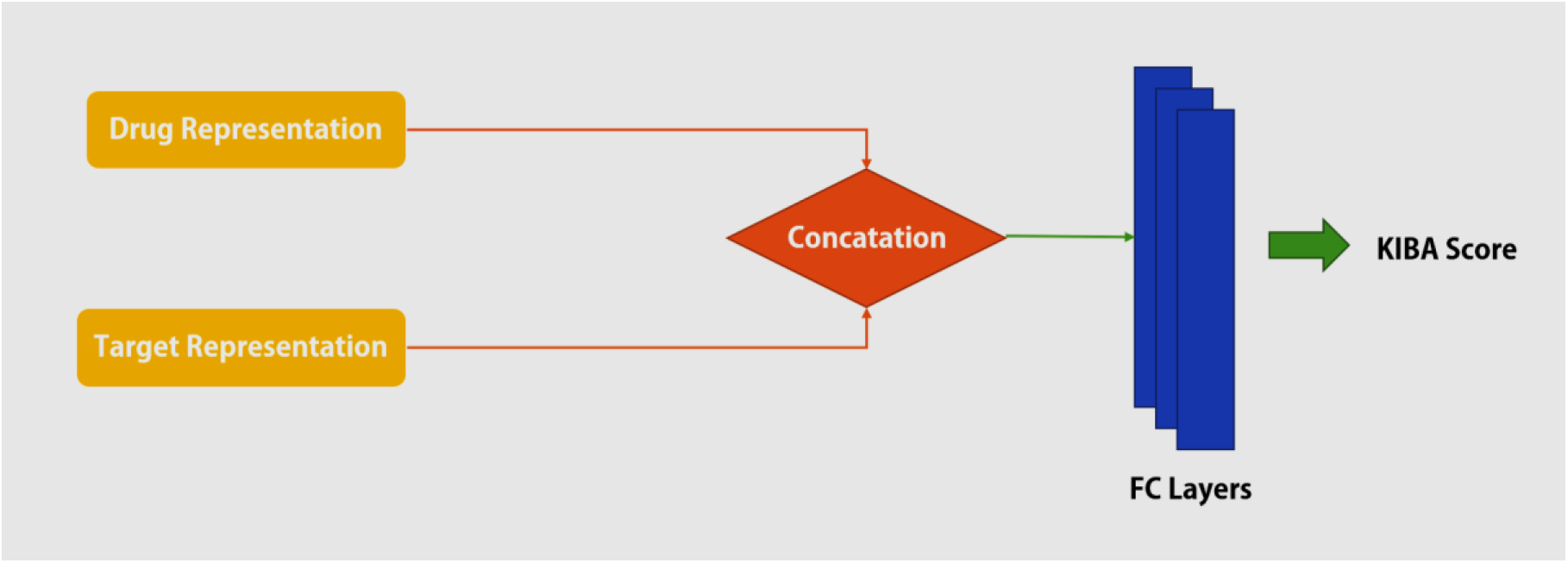
Step by step of our framework.

Different drug and protein representations are shown in the sets *D* and *P*, respectively as follows:

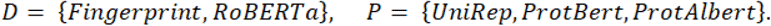

We evaluate the prediction results of our framework for each pair of the D×P set after training. So, we test six different models for the best linear representation for protein and drug. The number of nodes of fully connected layers varies depending on the size of the input representation vectors.

## III. Experiments and Results

We first describe the used dataset and the implementation details in this section. Then, we introduce used evaluation metrics to compare proposed models’ efficiency.

### A. Dataset

We evaluate our proposed models on KIBA[16] dataset, which has been popularly used as a benchmark for binding affinity prediction assessments[2], [5], [7]. The KIBA dataset is one of the large scale biochemical selectivity assays of the kinase inhibitors and originated from an approach called KIBA, in which kinase inhibitor bioactivities from different sources such as *K_i_*, *K_i_*, and *IC_50_* were combined[16]. In the KIBA dataset, the lower KIBA score corresponds to the higher binding affinity.

The original KIBA dataset contains a matrix of 467 proteins, and 52,498 drugs, with 246,088 interactions[16]. We use the filtered version of the KIBA dataset, in which each protein and ligand has at least ten interactions[5]. As a result, this dataset includes 229 unique proteins, 2,111 unique drugs, and 118,254 interactions.

Similar to DeepDTA[7], the KIBA scores pre-processed as follows: (i) negative of each KIBA score was taken, (ii) the minimum value among the negatives was chosen, and (iii) the absolute value of the minimum was added to all negative scores; thus the final form of the KIBA scores are constructed[7] and its value ranges from 0.0 to 17.2.

For drugs, we use Pubchem[15] to represent the molecular fingerprint and it contains data for only 2065 KIBA[16] drugs. In obtaining the representation of proteins, ProtAlbert[12] and ProtBert[12] need more than 32 gigabytes of RAM to run on protein sequences longer than 1000, which is not available to us. So, we consider protein sequences with a length of less than 1000 in the dataset, which is 185 proteins.

Due to limited resources, we consider a sample set of 1,512 interactions to train and test the models. We use 20% of the data for testing and the rest for training. More details of the sample set are shown in Table I.

**Table I.** Details of the sample set.

| No. Drugs | No. Proteins | No. Interactions | Split | No. samples |
| --- | --- | --- | --- | --- |
| 38 | 163 | 1512 | Train | 1209 |
|  |  |  | Test | 303 |

The sample set of interactions is constructed by the random sampling method. We also used Cochran’s formula[24] to calculate the sample size (error rate=0.025, Z=1.96). Therefore, the sample set size is equal to 1,512. Fig. 3 shows the total data distribution in the dataset and the data distribution in the sample set.

**Fig. 3.**
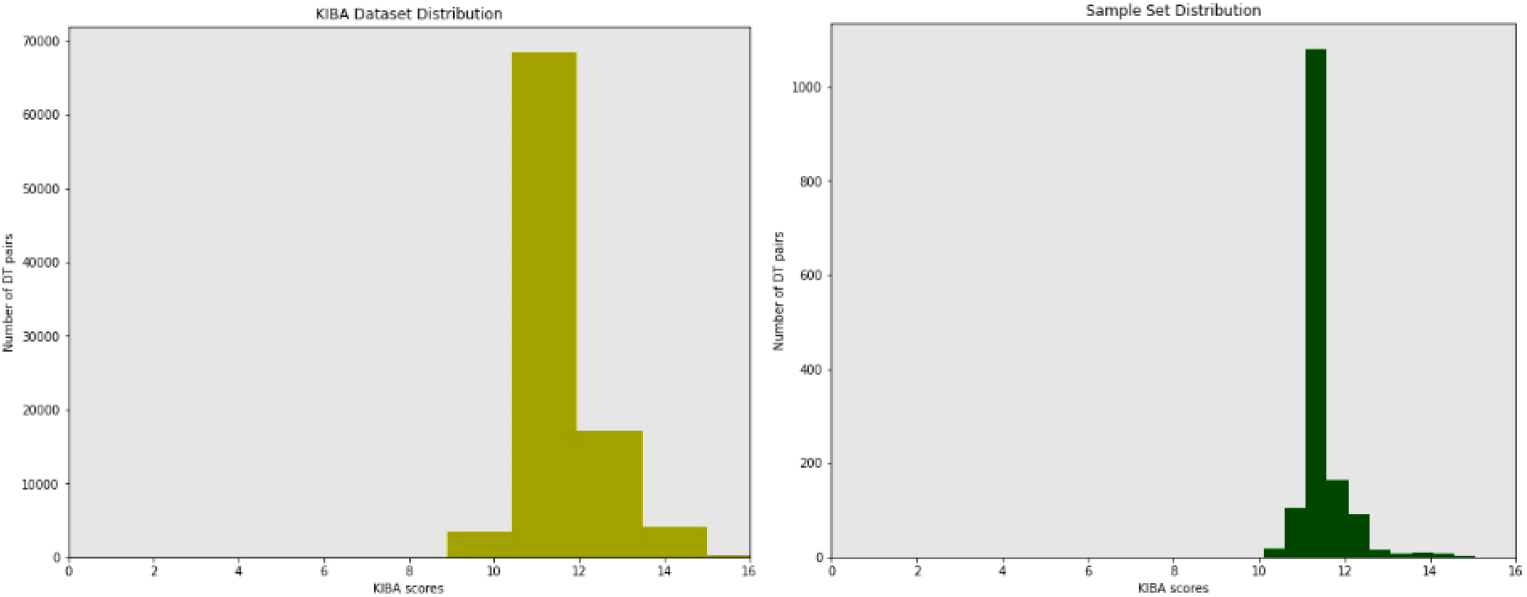
KIBA dataset and the sample set distributions.

This sample set is used to compare 6 models from ProtAlbert[21], ProtBert[21], and UniRep[20] models for protein representation and RoBERTA[22] and PubChem[23] fingerprint for drug representation.

### B. Implementation Details

We use the Python programming language and our experiments are run on Google Colaboratory[25]. The proposed framework is implemented using PyTorch[26] backend and ADAM optimization. We achieve a high performance of the proposed models with a relatively small range on hyperparameter tuning. The detailed settings are summarized in Table II.

**Table II.** Details of hyperparameter settings.

| Hyperparameter | Setting |
| --- | --- |
| Batch size | 1 |
| Learning rate | 0.003 |
| Optimizer | Adam |
| Activation function | Rectified Linear Unit (ReLU) |
| Loss function | Root Mean Square Error (RMSE) |

### C. Evaluation Metrics

This study performs our framework with different protein and drug representations. Then, the performance of our framework is compared to KronRLS[2], SimBoost[5], DeepDTA[7], GraphDTA[9], DGraphDTA[10] on KIBA[16] dataset. We use three metrics such as CI[27], MSE and RMSE to evaluate the performance in these models. In the following, we introduce these three metrics.

#### 1) Concordance Index (CI)

As suggested in [2], the CI can be used as an evaluation metric for the prediction accuracy in DTBA. CI[27] is a ranking metric for continuous values. The intuition behind the CI is whether the predicted binding affinity values of two random drug-target pairs were predicted in the same order as their actual values were or not. The CI ranges between 0.5 and 1.0, where 1.0 corresponds to perfect prediction accuracy and 0.5 corresponds to a random predictor. CI is computed as follows:

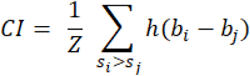

where *b_i_* is the prediction value for the larger affinity *s_i_*, *b_j_* is the prediction value for the smaller affinity *s_j_*, *Z* is a normalization constant that equals the number of data pairs with different label values, and *h*(*x*) is the Heaviside step function[2]. It is a discontinuous function and defined as:

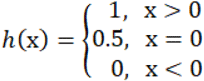

#### 2) Mean Square Error (MSE)

The MSE is a commonly used metric for the error in continuous prediction. It is used in regression tasks to measure how close the fitted line, represented by connecting the estimated values, is to the actual data points. Since DTBA is a regression task, we use the MSE as a metric:

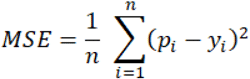

where *y_i_* is the actual output, *p_i_* corresponds to the prediction and *n* indicates the number of samples.

#### 3) Root Mean Square Error (RMSE)

RMSE is one of the regressor metrics. It is the distance, on average, of data points from the fitted line and computed as the square root of MSE:

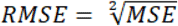

### D. Model evaluation

The experiments of this study are performed in two parts. First, the performance of different representations is compared with each other. Then, the model obtained from the best representations is compared with the state-of-art models[2], [5], [7], [9], [10].

#### 1) Representations Comparison

This study examines six different protein and drug representations on the sample set to find the best representation (Table III). We consider ProtAlbert[12], ProtBert[12] and UniRep[13] models for protein representation and RoBERTA[14] and PubChem[15] fingerprint for drug representation.

**Table III.** Prediction performance on a sample of KIBA dataset for different input representations, sorted by MSE.

| Drug Representation | Protein Representation | C-index | MSE | RMSE |
| --- | --- | --- | --- | --- |
| **FingerPrint** | **ProtAlbert** | **0.911** | **0.110** | **0.333** |
| FingerPrint | ProtBert | 0.885 | 0.115 | 0.339 |
| .FingerPrint | UniRep | 0.878 | 0.174 | 0.417 |
| .RoBERTa | UniRep | 0.879 | 0.216 | 0.465 |
| RoBERTa | ProtAlbert | 0.821 | 0.222 | 0.471 |
| RoBERTa | ProtBert | 0.884 | 0.249 | 0.499 |

Table III shows that molecular fingerprint representation of drugs has less MSE and more CI. So, it has performed better in DTBA prediction. This better performance could be because the component substructures are specified in the molecular fingerprint representation and the molecule’s structure is important in the binding affinity concept. Also, for linear representation of protein, the performance of transformers besides drug fingerprints is better than other representations. It shows that the ability of transformers to extract structural information from protein sequences is also helpful for DTBA. The best result of these six models is obtained using ProtAlbert[12] representation for the target protein and fingerprint for the drug. The architecture of this is shown in Fig. 4. We named this model TranDTA, which is a sequence-based model for predicting drug-target binding affinity.

**Fig. 4.**
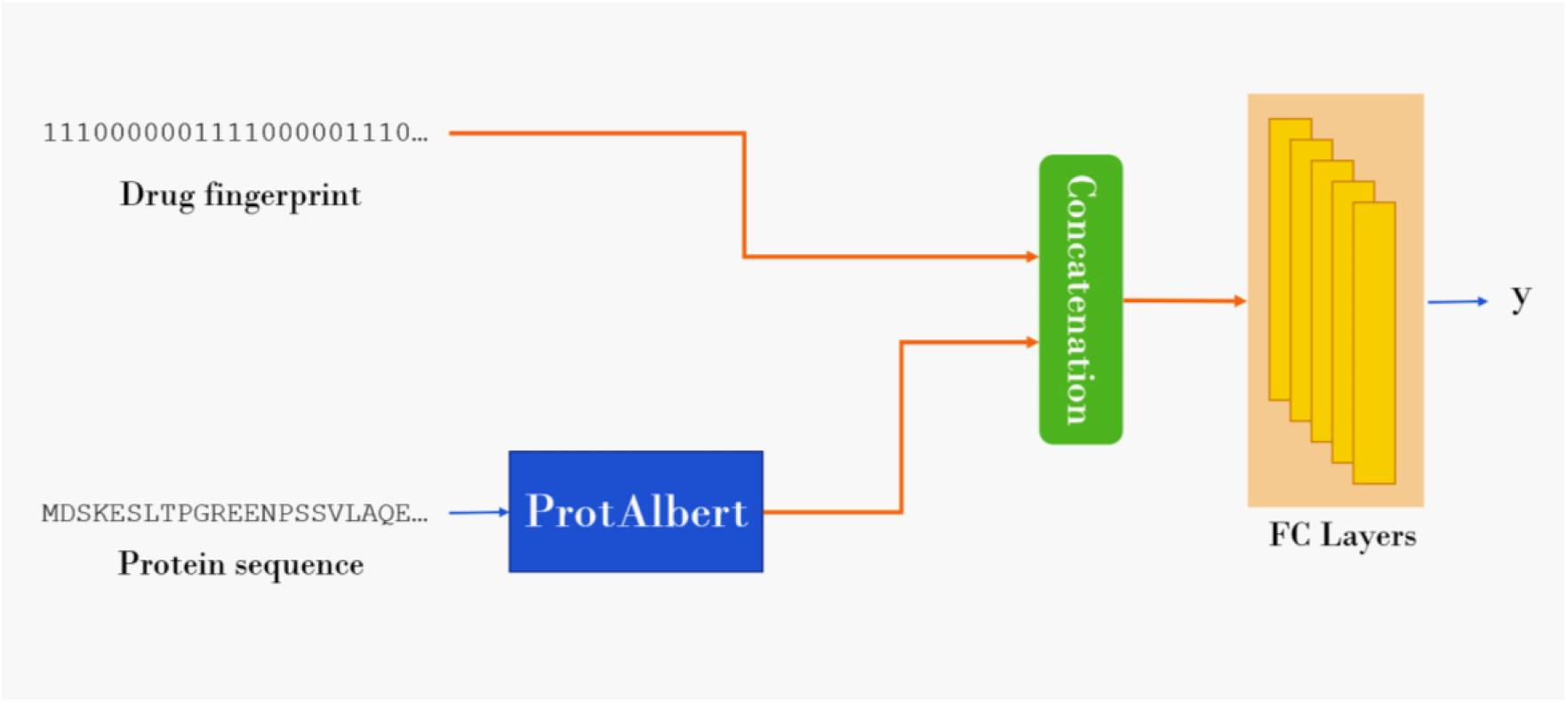
TranDTA architecture.

In TranDTA, we use a pre-trained model (ProtAlbert[12]) to transform the proteins sequences into feature vectors. Then, This vector is concatenated to the molecular fingerprint[19] vectors of drugs for feeding into five fully connected layers to predict the binding affinity value. We used 2048 nodes in the first FC layers. The following layers have 1024, 512, and 256 nodes, respectively.

#### 1) Comparison with state-of-art models

This study proposes a drug-target binding affinity prediction model, named TranDTA, based on only sequence information of drugs and proteins. Table IV shows the average CI, MSE, and RMSE values for KIBA[16] dataset.

**Table IV.** Prediction performance on KIBA dataset, sorted by MSE.

|  | DataSet | Model | CI | MSE | RMSE |
| --- | --- | --- | --- | --- | --- |
| Sequence-based | All | KronRLS | 0.782 | 0.411 | 0.620 |
|  | All | SimBoost | 0.836 | 0.222 | 0.471 |
|  | All | DeepDTA | 0.863 | 0.194 | 0.440 |
|  | Sample Set | TransDTA | **0.911** | **0.110** | **0.333** |
| Graph-based | All | GraphDTA | 0.891 | 0.139 | 0.372 |
|  | All | DGraphDTA | 0.904 | 0.126 | 0.354 |

The results show that the TranDTA performs much better than the similarity-based models (KronRLS[2] and SimBoost[5]) and the sequence-based models (DeepDTA[7]) in all three metrics. It also performs closely with the structure-based models that use graphs (GraphDTA[9] and DGrphDTA[10]) and is slightly better in this experiment.

Our model is competitive not only with sequence-based models[7] but also with models that use structural information[9], [10]. In addition, in our experience, TranDTA costs less time and calculations than other models. This reduction in time and calculations is because TranDTA does not learn input representations when the model is training; Instead, TranDTA uses pre-trained models to represent the protein. Therefore, this model can be trained and tested without limitations on resources (memory, CPU, and GPU).

## IV. Conclusion

So far, computational drug-target binding affinity prediction models have been extracting features using deep learning approaches. First sequence-based models and then graph-based models were introduced. Although graph-based models perform better in predicting, they require more resources to execute. Also, deep learning models that have been introduced so far learn the representations of inputs during training. Significantly, the sequence-based models, such as DeepDTA, limit the input sequences’ length and consider a fixed length that causes some information loss.

The proposed models in this study use the sequence of molecules to predict DTBA. We perform a study on pre-trained linear representations for protein and drug representations. New types of transformers are pre-trained on proteins and can use sequences to show structural features on their heads and layers. So, We assess protein sequence representation with ProtAlbert, ProtBert and UniRep models and drug representation with RoBERTA and PubChem fingerprint. We concatenate two extracted vectors from the model representations of the protein and drug s to feed a multi-layer fully connected neural network to predict DTBA. The Models are performed on KIBA, a benchmark dataset for binding affinity prediction evaluation. The results are competitive with other models that use the three-dimensional structure and representation of molecules to solve the DTBA problem. The increase in speed, decrease in calculations and the need for resources in the proposed models are due to two things:

a. Using sequences at the input of models instead of using two-dimensional and three-dimensional structures and representations
b. Using pre-trained models to represent molecular sequences means that there is no need to train the model to learn the representation of inputs in the neural network training section

On the other hand, the use of UniRep, ProtAlbert, ProtBert, and RoBERTa representations does not need to limit the length of sequences. It can take into account all the information in a sequence.

This paper proposes the novel method, named TranDTA, to predict DTBA. To the best of our knowledge, TranDTA is the first model that applies transformers to extract features of protein sequence and uses transformer representations in DTBA prediction. Experimental results show that TranDTA outperforms other sequence-based methods on the KIBA dataset prediction performance. Moreover, it performed closely with the structure-based models and was slightly better in this experiment.

Because of the success of transformers in natural language processing and the results of this study, we believe that TranDTA is a practical approach for DTBA prediction and can be pretty helpful in drug discovery development process.

There is a need to fine-tune TranDTA on the dataset for future work. In addition, a specific transformer can be used to represent the drug features to improve TranDTA,

